# Incidence, predictors and reasons for initial regimen modifications in patients on antiretroviral therapy in Witbank, South Africa, 2003-2017

**DOI:** 10.1101/610956

**Authors:** Moshibudi Poncho Bapela, Lazarus Rugare Kuonza, Alfred Musekiwa, Robert Summers

## Abstract

**Background:** Antiretroviral therapy (ART) is associated with unpleasant adverse effects that may require modification of regimens. ART modifications may lead to poor treatment outcomes. We determined the incidence, reasons and predictors for modification of initial ART regimens.

**Methods:** We retrospectively analysed data from Witbank pharmacovigilance sentinel site, South Africa. Censoring targeted the first incident of ART modification from the initial regimen. We included human immunodeficiency virus (HIV)-infected patients on ART, aged more than 18 years. We used the Cox-proportional hazard model to identify predictors for changing initial ART regimens.

**Results:** Among 2, 045 eligible patients, 38% (n=783) had their initial ART regimens changed. The overall incidence rate of ART modification was 10.0 per 100 person-years within a follow-up period of 7 794.6 person-years (PYs). Reasons for changing were adverse drug reactions (ADRs) (60%), prescriber’s decisions (37%), drug toxicity (26%) and treatment failure (12%). The most commonly changed regimens were stavudine (68%) and zidovudine (44%) based regimens. Stavudine-based regimen had the highest changing rate of 13.6 per 100 PYs compared to zidovudine (8.0 per 100 PYs) and tenofovir (6.5 per 100 PYs). Using tenofovir as reference, stavudine (aHR 2.3; 95% CI 1.8-2.9; p<0.001) and zidovudine (aHR 1.5; 95% CI 1.2-3.2; p<0.001) based regimens were significantly associated with regimen modifications. The predictors for changing ART regimens included drug toxicity (aHR 2.6; 95% CI 2.1-3.1), ADRs (aHR 2.1; 95% CI 1.3-3.2), treatment failure (aHR 2.0; 95% CI 1.5-2.4), baseline cd4 count of ≥200 (aHR 1.7; 95% CI 1.3-2.1) and initiation regimens (stavudine and zidovudine).

**Conclusion:** The findings were suggestive of a moderate incidence of initial ART regimen changing. Patients on stavudine and zidovudine based regimens changed primarily due to ADRs and drug toxicity. We recommended that clinicians should consider changing patients who are still on stavudine-containing regimens, however, changing should be individualized.

## Introduction

Human immunodeficiency virus (HIV) infection is a retroviral infection that causes acquired immunodeficiency syndrome (AIDS), a spectrum of conditions resulting from supressed immunity [1, 2]. Though HIV is the causative agent for AIDS, most of the morbidity and mortality related to AIDS is associated with opportunistic infections and tumours that are uncommon in HIV-uninfected people [2-3]. HIV continues to be a major challenge for public health globally [4-6]. The World Health Organization (WHO) reported that 36, 9 million people were living with HIV worldwide by the end of 2017 [4-5].

South Africa (SA) continues to be the country with the highest number of HIV-infected people globally. The prevalence of HIV in SA was 7.1 million by the end of 2017, a major increase from 4.72 million in 2002 [4, 7-8]. However, new infections declined yearly from 1.77% in 2002 to 1.27% in 2016 [7-8]. The country has the largest antiretroviral programme worldwide, which contributed to reduction in AIDS-related morbidity and mortality [6, 8-9]. Antiretroviral treatment/therapy (ART) consists of drugs used in HIV medicine to reduce the complications arising from HIV-related illnesses but do not cure the condition [2].

The treatment has changed HIV from a life-threatening to a manageable chronic condition over the years [3, 10-13]. ART not only prolongs life, but also significantly limits transmission to those who were HIV-uninfected [1, 6, 10, 14-15]. Due to complexity of HIV, multiple combinations of antiretroviral drugs (long-term) are required to control its replication optimally and manage costs [10-11, 16-17]. The drugs are often associated with unpleasant adverse effects that may require early modification of regimens [1, 16-18]. Treatment regimen switch and discontinuation may precipitate subsequent development of drug resistance [1, 16-18].

The major drivers for regimen modifications world-wide included treatment failure and adverse drug reactions (ADRs) [17-20]. With improvement in ART over years, the factors associated with modifying regimens changed from merely treatment failure and drug resistance to several factors like non-adherence, drug interactions and drug related toxicity [14]. However, drug toxicity still appears to be the main reason for changing ART regimens. Most toxicity was found with stavudine (d4T) and zidovudine (AZT) containing regimens [3, 12]. The regimens can be switched for convenience, tolerability, simplification of treatment, potential new drug interactions, pregnancy and plans for pregnancy, elimination of food restrictions, virological failure and drug toxicities [1, 8-9, 11, 18].

Patients on d4T, didanosine or AZT-containing regimens may also have been changed even when they were doing well and were satisfied with their treatment to prevent the long-term toxicity associated with these drugs [3, 13]. The factors to be considered before a regimen is modified, include drug resistance, potential adverse events, drug interactions, diet and cost [10, 12, 14]. In resource poor settings, a number of patients remain on failing first-line regimens due to limited treatment options, poor laboratory facilities, lack of confidence in health care providers making switches and delay in changing [21].

Other studies conducted in South Africa, have reported higher rates of treatment modification in stavudine and zidovudine containing regimens compared to tenofovir-containing regimens [20, 22]. In order to ensure antiretroviral drug safety, it is essential to understand drug effects, prescribing patterns, and regimen modification patterns (and reasons). Identification of risk factors for early treatment modification is essential to inform/guide treatment policies and guidelines. The aim of the study was to determine the incidence of, predictors and reasons for changing of initial antiretroviral treatment regimens.

## Methods

### Study design and setting

We conducted a retrospective cohort study which entailed analysis of secondary data collected from the Witbank sentinel treatment site, located in Mpumalanga Province of South Africa. This site is situated on the Highveld of Mpumalanga Province, in Nkangala district municipality. It is an expanding urban area, surrounded by coal mining industries with 22 coal mines, several power stations and steel mills. With the increase in mining activities, pollution becomes a major problem with many wetlands, streams and rivers now obstructed with coal dust. Witbank clinic caters for clients from all over Witbank area (people staying in central business district, those working near the clinic and patients from different areas who do not want to utilize nearest clinics due to stigma). Witbank clinic is a comprehensive primary health care centre which offers all the primary health care services. The clinic provides treatment for common illnesses, long term disorders (HIV, diabetes mellitus, hypertension and mental health) and prevention of future illnesses through health education, immunizations and screening programmes. The clinic enrols HIV-infected patients, initiate ART and change ART regimens following the national guidelines. Clinical and laboratory evaluations were done before ART initiation. The clinic provided ART to patients on a monthly basis. Cotrimoxazole was provided to all the patients with the CD4 cell count of <200 cells/mm3. The clinic followed the national guidelines for recommended first-line ART regimens. The standardised first-line ART regimen consisted of two nucleoside reverse transcriptase inhibitors (NRTI) and one non-nucleoside transcriptase inhibitors (NNRTI). Modifications/changing of ART regimens are done following the prescribed guidelines and standardised procedures. Adverse drug reactions were recorded on the standardised form in cases of drug substitutions due to intolerance. In 2015, pharmacovigilance programme which was initially operating from Witbank Hospital, moved to Witbank clinic. Since its inception, the programme enrolled 2, 216 HIV-infected patients who were already on ART. The programme monitors adverse drug reactions of antiretroviral drugs and related medicines.

### Study population

The study population consisted of HIV-infected patients on ART enrolled in a sentinel treatment site for the Pharmacovigilance Research Centre based at Sefako Makgatho Health Sciences University (SMU), previously the Medical University of Southern Africa (MEDUNSA). We included only patients older than 18 years, initiated antiretroviral therapy between March 2003 and December 2017. The study excluded patients less than 19 years of age, those without recorded initiation date, missing regimen on initiation, registered in 2018 and ART changing date earlier than initiation date. The study population was followed up from the initiation date. No sample size was calculated as the entire cohort from March 2003 to December 2017 was investigated.

### Data collection

We extracted data from the SMU Pharmacovigilance Centre database. Patients were enrolled at the treatment site by an onsite coordinator from March 2007. The coordinator engaged with patients by interviewing them after obtaining their informed consent (for participation in the pharmacovigilance survey). Enrolling new patients involved retrospective collection of data from the day of ART initiation till enrolment date. The first ART initiation record dated March 2003. Files of enrolled patients were marked with the enrolment date on the cover. Case report forms (CRFs) were completed on each visit by interviewing the patients and consulting their medical records. The CRF covered demographic, life style, clinical, pharmacy (ARV treatment and other medication for the patient), laboratory test results and patient status. The information was captured on Pharmacovigilance Surveillance System Version 1.0.0 at the Research Centre. The data were then exported into a spreadsheet database (wide format) and analysed.

### Variables

The primary outcome was first-time ART modification after initiation of treatment. The modification included changing patient treatment to an alternative first-line or second-line regimen. Patients may also have been changed to non-standardised ART treatment. The initiation date was considered as the baseline. The end date of the failure event was the initial ART regimen changing date. Censoring targeted the initial ART regimen changing date, loss to follow-up before event, died before developing the event and those who reached the end of the study without the event. We collected independent variables that included demographics (age, sex, race); smoking and alcohol consumption history; clinical and laboratory data (chronic and opportunistic illnesses, WHO stage at initiation, baseline CD4 count, baseline viral load, haemoglobin and body mass index), ADRs and treatment variables (initiation date, regimen changing dates and initiation regimen).

### Definitions

We defined treatment discontinuation as stopping any antiretroviral drug or regimen temporarily or permanently due to toxicity. Regimen changing/modification/switching was defined as changing the ART regimen from one standardised regimen to another or to a non-standardised (individualised) regimen. Substitution of a drug was defined as replacement of one or more drugs within the regimen either standardised or non-standardised to another more potent drug.

### Statistical Management

Data cleaning was done using Microsoft Excel and STATA. The data were de-identified to ensure confidentiality and analysed using STATA statistical software (version 15, StataCorp, College Station, TX, USA).

### Data analysis

We described categorical variables using proportions and frequencies and compared them using Pearson’s chi square test. We calculated the frequency of initial ART modification and related characteristics. We calculated proportions to determine the frequency of adverse effects of different treatment regimens, and related characteristics. The Fisher’s exact test was used to test for statistical significance of differences in frequency of ADRs between drug regimens. We used Cox proportional hazard model to determine predictors for changing ART regimens. We used a manual forward step-wise method to identify the independent predictor variables. The p-value of 0.2 was used as a cut-off value in bivariate analyses in order to include the variables in the multivariate model. The variables were entered in the multivariate model by starting with the one with the smallest p-value. We considered the p value ≤ 0.05 as statistically significant in the multivariate analysis. The rates of ART modification were calculated per 100 patient-years with 95% confidence intervals. We used the log-rank test to compare the survival distributions of different treatment regimens. Kaplan-Meier (KM) survival analysis was used to measure the cumulative probability of regimen modification over time by treatment regimens and adverse drug reactions. We utilised Kaplan-Meier survival curves to display the findings. We used frequency tables and graphical displays to summarise and display data.

## Ethical considerations

We excluded personal identifying variables during analysis to ensure anonymity. Permission to conduct the study at the facility was granted by the Provincial Department of Health, Mpumalanga Province. The study was approved by the Faculty of Health Sciences Research Ethics Committee of the University of Pretoria (reference number 462/2017).

## Results

A total of 2,216 HIV-infected patients initiated on ART were enrolled in the pharmacovigilance programme from March 2007 to February 2018. However, the patients were retrospectively followed up from first initiation date (First record dated March 2003). We analysed data for 2,045 after excluding patients who were less than 19 years of age at initiation (n=50), had missing ART resumption date (n=5), missing ART regimens on initiation (n=17), changed ART before initiation (n=89), missing time to event (n=2) and those enrolled in 2018 (n=8) (Fig 1).

**Fig 1.**
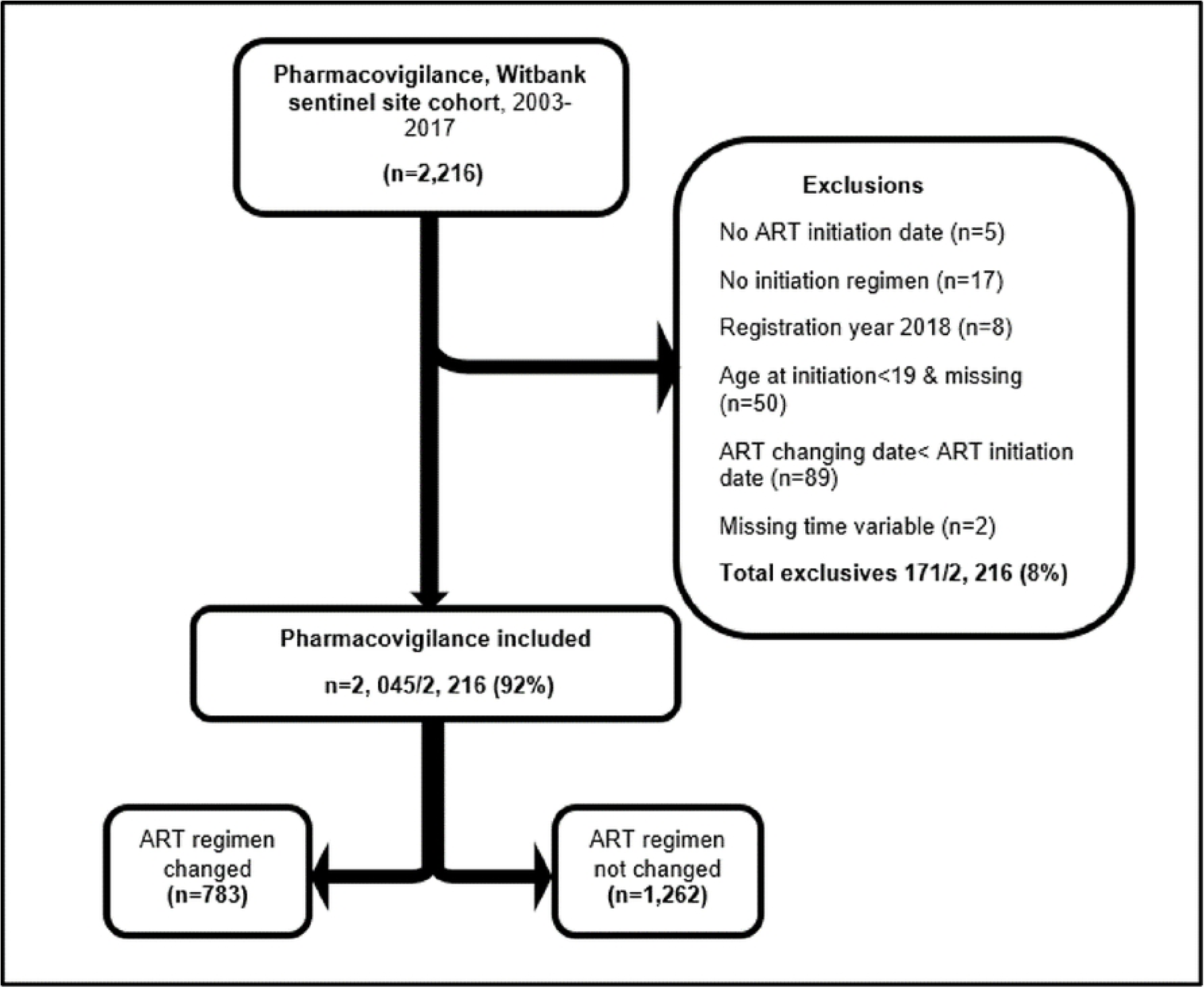
Flow chart indicating inclusion and exclusion criteria of HIV-infected patients on ART, Witbank, South Africa, March 2003-December 2017.

### Characteristics of the study population

Of the 2,045 eligible patients, 783 (38%) had their initial ART regimens changed (Fig 1). There were 4% (n=83) of patients who had individual drug dosage substitution, 63% (n=52) had their regimens changed. Other patients (8%, n=158) had their antiretroviral treatment discontinued temporarily. The median age at ART initiation was 36 years (IQR 30-44 years) and 65% (n=1,325) of the patients were female. Majority of the patients were black (99.8%). Approximately half of the patients were initiated on TDF-based regimens (51%, n=1,054), 35% (n=718) on d4T-based and 10% (n=212) on AZT-based regimens. Some patients were initiated on non-standardised regimens (3%, n=61) (Table 1). The age group with the highest frequency was 25-34 years (38%), followed by 35-44 years (32%). These age groups had higher frequencies in all ART initiation regimens. Female patients predominated in all ART regimens. Among patients with recorded CD4 cell count, 59% (n= 1,208) were initiated with the CD4 count of ≥200 cells per cubic millimetre (cells/mm3). Majority of the patients had no baseline viral load (VL) results 88% (n=1,797) and WHO stages 68% (n=1,397). Of those with available results, 10% had viral loads below 1,000 copies/l. Thirty-one percent were initiated at WHO-stages 1 or 2. Chronic illnesses, opportunistic infections and the use of traditional medicines was recorded for less than 3% of patients. Nine percent (9%) of patients consumed alcohol and five percent smoked cigarette. Sixty-three percent (n=1,298) of patients used other concomitant drugs (Table 1).

**Table 1.** Baseline sociodemographic and clinical characteristics of HIV-infected patients on ART, Witbank, South Africa, 2003-2017

| **Variable** |  | **Overall (n=2,045), n (%)** |
| --- | --- | --- |
| Age categories | 19-24 | 144 (7.0) |
|  | 25-34 | 785 (38.4) |
|  | 35-44 | 674 (33.0) |
|  | 45-54 | 348 (17.0) |
|  | 55+ | 94 (4.6) |
| Sex | Female | 1 325 (64.8) |
|  | Male | 720 (35.2) |
| Race | Black | 2 040 (99.8) |
|  | Coloured | 5 (0.2) |
| WHO stage at initiation | I/II | 636 (31.1) |
|  | III/IV | 12 (0.6) |
|  | Missing | 1,397 (68.3) |
| CD4 count at initiation | <200 | 463 (22.6) |
|  | ≥200 | 1,208 (59.1) |
|  | Missing | 374 (18.3) |
| Viral load at initiation | <1000 | 185 (9.1) |
|  | ≥1000 | 29 (1.4) |
|  | Missing | 1,831 (89.5) |
| Cigarette smoking | Yes | 106 (5.2) |
|  | No | 1,938 (94.8) |
| Alcohol consumption | Yes | 192 (9.4) |
|  | No | 1,852 (90.6) |
| Adverse drug reactions | Yes | 551 (27.0) |
|  | No | 1,494 (73.0) |
| Pregnancy | Yes | 69 (3.4) |
|  | No | 1,976 (96.6) |
| Concomitant drugs<sup>a</sup> | Yes | 1,298 (63.5) |
|  | No | 747 (36.5) |
| Traditional medicine use | Yes | 14 (0.7) |
|  | No | 2,031 (99.3) |
| Chronic illnesses<sup>a</sup> | Yes | 32 (1.6) |
|  | No | 322 (15.7) |
|  | Missing | 1,691 (82.7) |
| Opportunistic infections<sup>a</sup> | Yes | 42 (2.1) |
|  | No | 308 (15.1) |
|  | Missing | 1,695 (82.8) |
| NRTI backbone | d4T-based | 718 (35.1) |
|  | AZT-based | 212 (10.4) |
|  | TDF-based | 1,054 (51.5) |
|  | Non-standardised | 61 (3.0) |

Abbreviations: NRTI, Nucleoside Reverse Transcriptase Inhibitors. aOpportunistic infections: tuberculosis, wasting syndrome, cervical cancer; aChronic illnesses: Diabetes Mellitus, Epilepsy, Hypertension; a Concomitant drug: any other drug taken excluding ART

### Treatment changes

The frequencies of regimen changing for d4T-based, AZT-based, and non-standardised regimens were 68% (n=490), 44% (n=93), and 34% (n=21), respectively. Majority of patients were on TDF-based regimens, however, the frequency of changing (17%, n=179) was lower than all other regimens. Females accounted for 66% (n=521) of the patients who had their initial ART regimens changed. Half (n=34/68) of the pregnant female patients had their treatment changed. Of those who had chronic diseases and opportunistic infections, over 70% had their treatment changed. Twenty-seven percent (n=551) of the patients experienced ADRs. Of the patients with ADRs, 60% (n=329) had their initial ART regimens changed (Table 2). There were 8% (n=158) who had their treatment discontinued temporarily. Forty-one percent (n=64) of those who had their ART discontinued temporarily also had their regimens changed.

**Table 2.** Frequency of regimen changes and related characteristics, Witbank, South Africa, March 2003-December 2017

| Variable |  | Changed regimen<br>(n=783), n (%) | Sustained regimen<br>(n=1,262), n (%) | p-value |
| --- | --- | --- | --- | --- |
| Age at initiation | 19-24 | 45 (31.2) | 99 (68.7) | 0.081 |
|  | 25-34 | 320 (40.8) | 465 (59.2) |  |
|  | 35-44 | 266 (39.5) | 408 (60.5) |  |
|  | 45-54 | 120 (34.5) | 228 (65.5) |  |
|  | 55+ | 32 (34.0) | 62 (66.0) |  |
| Sex | Female | 521 (39.3) | 804 (60.7) | 0.193 |
|  | Male | 262 (36.4) | 458 (63.6) |  |
| WHO stage at initiation | I/II | 153 (24.1) | 483 (75.9) | <0.001 |
|  | II/IV | 2 (16.7) | 10 (83.3) |  |
|  | Missing | 628 (45.0) | 769 (55.0) |  |
| CD4 count at initiation | <200 | 96 (20.7) | 367 (79.3) | <0.001 |
|  | ≥200 | 571 (47.3) | 637 (52.7) |  |
|  | Missing | 116 (31.0) | 258 (69.0) |  |
| Viral load at initiation | <1000 | 3 (30.0) | 7 (70.0) | 0.404 |
|  | >1000 | 13 (44.8) | 16 (55.1) |  |
|  | Missing | 709 (38.7) | 1,122 (61.3) |  |
| Pregnancy | Yes | 34 (50.0) | 34 (50.0) | 0.096 |
|  | No | 749 (37.9) | 1, 227 (62.1) |  |
| Cigarette smoking | Yes | 45 (42.4) | 61 (57.6) | 0.488 |
|  | No | 738 (38.1) | 1,200 (61.9) |  |
| Alcohol consumption | Yes | 66 (34.4) | 126 (65.6) | 0.367 |
|  | No | 717 (38.7) | 1,135 (61.3) |  |
| Adverse drug reactions | Yes | 329 (59.7) | 222 (40.3) | **<0.001** |
|  | No | 454 (30.4) | 1, 040 (69.6) |  |
| Concomitant medicines | Yes | 526 (40.5) | 772 (59.5) | **0.006** |
|  | No | 257 (34.4) | 490 (65.6) |  |
| Chronic illnesses | Yes | 23 (71.9) | 9 (28.1) | **<0.001** |
|  | No | 192 (59.6) | 130 (40.4) |  |
|  | Missing | 568 (71.9) | 1,123 (28.1) |  |
| Traditional medicines | Yes | 9 (64.3) | 5 (35.7) | **0.045** |
|  | No | 774 (38.1) | 1,257 (61.9) |  |
| Opportunistic infections | Yes | 32 (76.2) | 10 (23.8) | **<0.001** |
|  | No | 180 (58.4) | 128 (41.6) |  |
|  | Missing | 571 (33.7) | 1,124 (66.3) |  |
| Drug toxicity | Yes | 214 (88.4) | 28 (11.6) | **<0.001** |
|  | No | 569 (31.6) | 1,234 (68.4) |  |
| Body mass index | <18.5 | 12 (24.0) | 38 (76.0) | 0.124 |
|  | 18.5-24.9 | 157 (32.0) | 334 (68.0) |  |
|  | 25-29.9 | 107 (30.2) | 247 (69.8) |  |
|  | 30+ | 272 (34.5) | 516 (65.5) |  |
|  | Missing | 5 (62.5) | 3 (37.5) |  |
| Treatment failure | Yes | 115 (75.7) | 37 (24.3) | **<0.001** |
|  | No | 668 (35.3) | 1,225 (64.7) |  |
| NRTIs | Stavudine | 490 (68.2) | 228 (31.7) | **<0.001** |
|  | Zidovudine | 93 (43.9) | 119 (56.1) |  |
|  | Tenofovir | 179 (17.0) | 875 (83.0) |  |
|  | Non-standardised | 21 (34.4) | 40 (65.6) |  |

The overall incidence rate of initial ART regimen changes was 10.0 per 100 person-years (PYs) (95% Confidence Interval (CI): 9.4-10.8) over a total follow-up period of 7794.6 PYs. Of the 2,045 included in the study, 0.6% (n=12) died, 0.9% (n=18) were lost to follow-up (LTFU) and 61% (n=1,246) reached the end of the study without developing the event (ART regimen changing). Median times to death, LTFU and end of study before ART regimen changing were 3 years (IQR:1.8-4.2 years), 2 years (IQR: 1.3-3.0), and 3 years (IQR:1.7-7.8), respectively. Using a non-standardised regimen as reference, cumulative PYs of exposure was shortest for AZT-based (1155.7 PYs) compared to TDF-based (2747.5 PYs) and d4T-based (3600.1 PYs). The rates of changing ART regimens were higher in patients initiated on d4T (13.6; 95% CI 12.5-14.9), AZT (8.0; 95%CI 6.6-10.0), non-standardised (7.2; 95% CI 4.7-11.0) as compared to TDF (6.5; 95% CI 5.6-7.5) per 100 PYs.

Kaplan-Meier (KM) survival plots for the length of time after ART initiation until occurrence of the first regimen change were presented for the different backbone NRTIs groups (Fig 2). There was a significant difference in survival times between the backbone NRTIs groups (log rank test=0.001). Patients on TDF-based regimens had a better survival on the initial ART regimen, than those on d4T and AZT based regimens. Patients on d4T based regimens had the shortest survival time.

**Fig 2.**
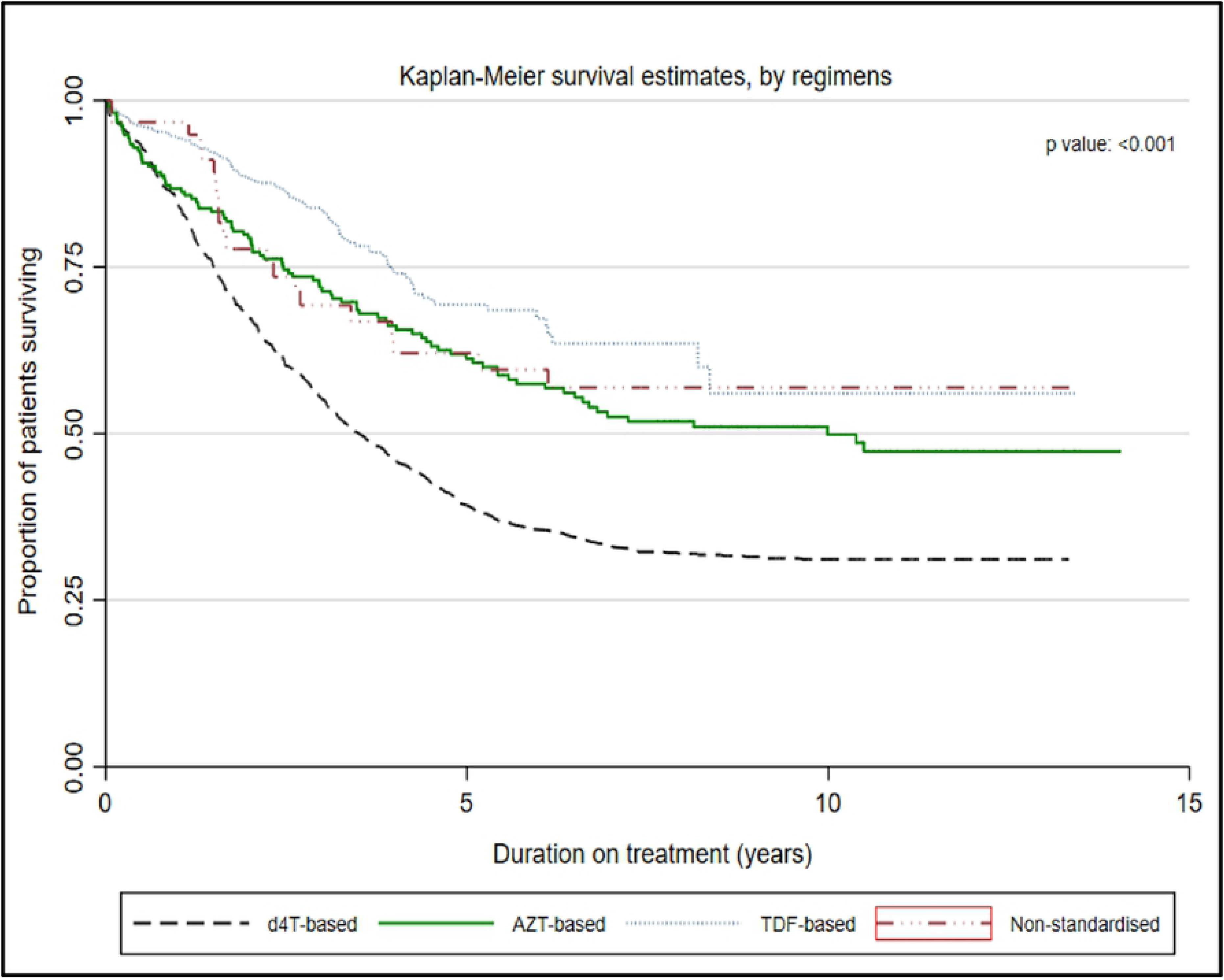
Kaplan-Meier analyses of time to first ART regimen change by backbone NRTI regimens among HIV-infected patients, Witbank, South Africa, March 2003-December 2017.

### Reasons for changing ART

The main reason for initial ART regimen changing was ADRs 60%, followed by prescriber’s decision 37%, drug toxicity 26% and treatment failure 15%. Stavudine-based regimens accounted for the majority of ART regimen changes due to toxicity (86%, n=176) and ADRs (81%, n=267). Approximately half of the patients with treatment failure (58%) were initiated on d4T-based regimens. Among the pregnant women who had their regimens changed, 59% (n=20) were on d4T-based regimens. Patients on tenofovir-based regimens were changed from a non-combined drugs to a fixed-dose combination (FDC) pill. Guidelines related (7.5%) and unavailability of ART (2.6%) were the other reasons for ART regimen changing (Table 3).

**Table 3.** Reasons for ART regimen changing according to classes of regimens for HIV-infected patients, Witbank, South Africa, March 2003-December 2017

| Reasons for changing | Overall ART changes, n | D4T-based n (%) | AZT-based n (%) | TDF-based n (%) | Non-standardised n (%) |
| --- | --- | --- | --- | --- | --- |
| ADRs <sup>a</sup> | 329 | 267 (81.2) | 28 (8.5) | 27 (8.2) | 7 (2.1) |
| Prescriber decision | 289 | 113 (39.1) | 51 (17.6) | 114 (39.4) | 11 (3.8) |
| Toxicity | 204 | 176 (86.3) | 14 (6.9) | 13 (6.4) | 1 (0.5) |
| Treatment failure | 115 | 67 (58.3) | 16 (13.9) | 30 (26.1) | 2 (1.7) |
| Guidelines | 59 | 51 (86.4) | 2 (3.4) | 3 (5.1) | 3 (5.1) |
| ARV <sup>a</sup> unavailable | 20 | 12 (60.0) | 2 (10.0) | 6 (30.0) | 0 (0) |
| Pregnancy | 34 | 20 (58.8) | 7 (20.6) | 5 (14.7) | 2 (5.9) |
| Drug interaction | 5 | 3 (60.0) | 2 (40.0) | 0 (0) | 0 (0) |
| Poor adherence | 3 | 3 (100) | 0 (0) | 0 (0) | 0 (0) |
| Unknown | 59 | 47 (79.6) | 5 (8.5) | 5 (8.5) | 2 (3.4) |

aADRs: adverse drug reactions; aARV: antiretroviral drugs;
\*Some patients had more than one reason for changing

### Adverse drug reactions

The overall prevalence rate of ADRs was 27% (n=551), with female patients predominating (72%, n=394). There was a significant difference between male and female patients on development of ADRs (p-value <0.001). Of those who experienced ADRs, 60% (n=329) had their regimens changed. Among the 158 patients who had their ART stopped temporarily, 57% (n=90) had ADRs and the majority (74%) were initiated on d4T-based regimens. Patients on d4T-based regimens had higher frequency of ADRs 72% (n=399), followed by AZT-based regimens 13% (n=74) and TDF-based 10% (n=60). There was a significant difference between initial ART regimens in developing ADRs (p<0.001). The commonly experienced ADRs were lipodystrophy, peripheral neuropathy, pain, neurological problems and rash accounting for 43% (n=235), 34% (n=189), 22% (n=124), 20% (n=109) and 14% (n=79) respectively (Table 4).

**Table 4.** Distribution of adverse drug reactions by ART regimens among HIV-infected patients, Witbank, South Africa, March 2003-December 2017

| Adverse drug reactions | ADRs (n=551) | D4T-based<br>n (%) | AZT-based<br>n (%) | TDF-based<br>n (%) | Non-standardised<br>n (%) |
| --- | --- | --- | --- | --- | --- |
| Lipodystrophy | 235 (42.6) | 180 (76.6) | 38 (15.3) | 14 (6.0) | 5 (2.1) |
| Peripheral neuropathy | 189 (34.3) | 143 (75.6) | 33 (17.5) | 9 (4.8) | 4 (2.1) |
| Pain | 124 (22.5) | 91 (73.4) | 21 (16.9) | 11 (8.9) | 1 (0.8) |
| Neurological problems | 109 (19.8) | 76 (69.7) | 11 (10.9) | 12 (11.0) | 8 (7.4) |
| Skin rash | 79 (14.3) | 60 (75.9) | 9 (11.4) | 9 (11.4) | 1 (1.3) |
| Gastrointestinal | 72 (13.2) | 49 (68.1) | 10 (13.9) | 10 (13.9) | 3 (4.2) |
| Breast issues | 18 (3.3) | 13 (72.2) | 1 (5.6) | 3 (16.7) | 1 (5.6) |
| Lethargy | 14 (2.5) | 10 (71.5) | 1 (7.1) | 2 (14.3) | 1 (7.1) |
| Skin discoloration | 8 (1.5) | 6 (75.0) | 1 (12.5) | 1 (12.5) | 0 (0) |

Kaplan-Meier plots of the probability of changing initial ART regimen show that patients who did not experience ADRs had better survival rates on the initial ART regimen than those who experienced ADRs (Fig 3).

**Fig 3.**
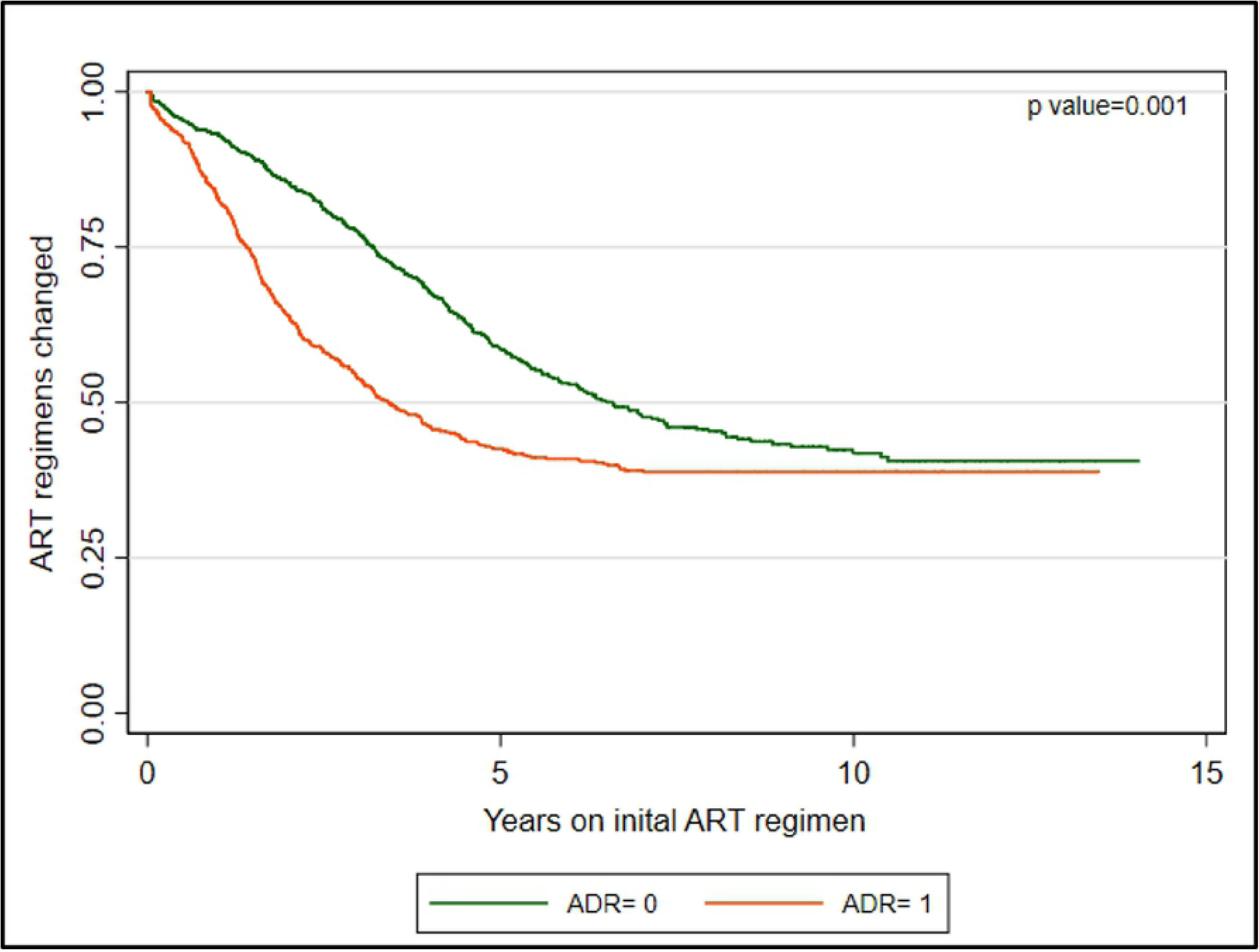
Kaplan Meier analyses of first ART regimen change due to ADRs among HIV-infected patients, Witbank, South Africa, March 2003-December 2017.

### Predictors of ART changing

There was evident increase in ART regimen changing attributable to ADRs, drug toxicity, treatment failure and prescribed initiation regimens. We analysed the time to the first incidence on initial ART regimen changing using Kaplan-Meier plots, stratified by backbone first-line antiretroviral regimens (Fig 2). Using TDF-based regimens as reference, the results from Cox proportional hazard analysis found patients using d4T-based (adjusted Hazard ratio (aHR): 2.3; 95% CI 1.8-2.9, p<0.001), AZT (aHR: 1.5; 95% CI 1.0-2.2, p=0.026) and non-standardised (aHR 1.6; 95%CI 0.9-2.9, p=0.113) regimens as significantly associated with regimen changing. Other predictors of ART regimen changing included drug toxicity (aHR 2.6; 95% CI 2.1-3.1, p<0.001), ADRs (aHR 2.1; 95%CI 1.3-3.2, p=0.001), treatment failure (aHR 1.9; 95%CI 1.6-2.4, p<0.001) and cd4 cells of 200 and more cells/mm3 (aHR 1.7; 95% CI 1.3-2.1, p<0.001). There was a significant interaction between ADRs and initiation regimens that was included in the Cox proportional hazard model (Table 5).

**Table 5.**
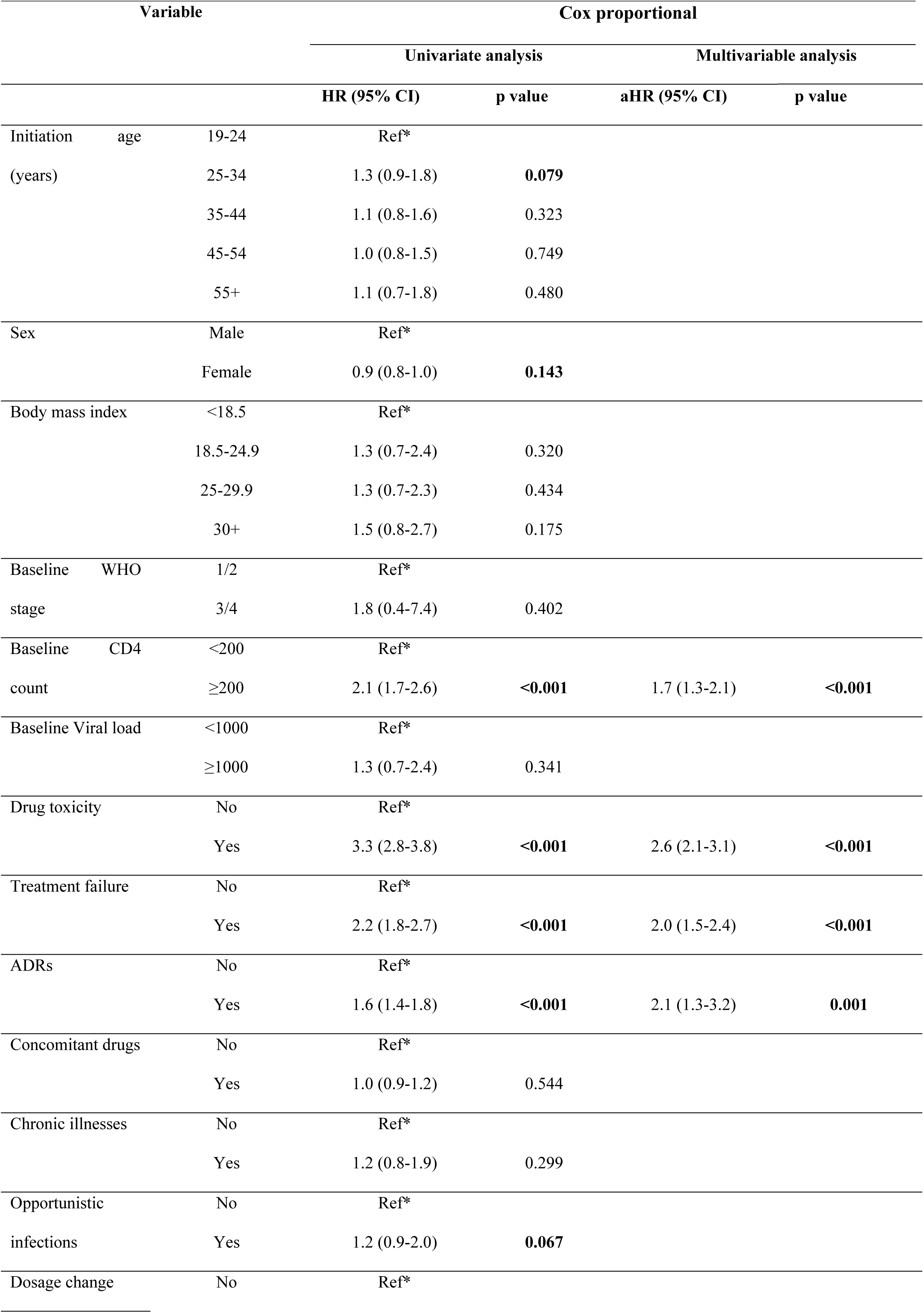
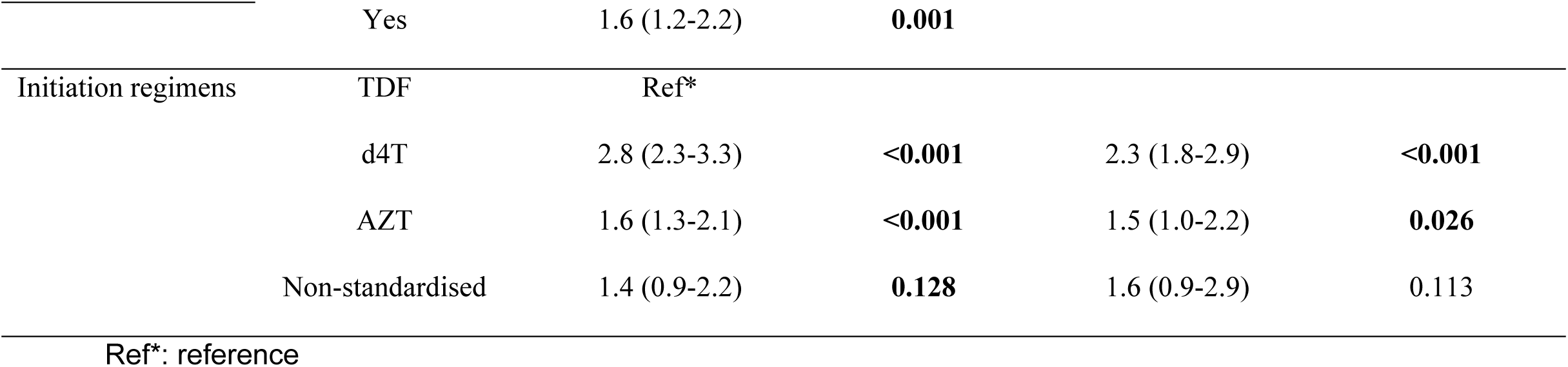
Cox proportional hazard model of predictors for initial ART regimen change among HIV-infected adults, Witbank, South Africa, March 2003-December 2017

## Discussion

This study reports the occurrence of and the reasons for initial ART regimen changing in a pharmacovigilance sentinel site over a 14-year period (2003-2017). Of the 2,045 patients who were on ART during a median follow-up of six years (IQR: 2.4-8.2 years), approximately one-third had their regimens modified. The specific incident rates of total regimen changing for patients on d4T, AZT, non-standardised and TDF based regimens were 13.6 per 100 PYs, 8.0 per 100 PYs, 7.2 per 100 PYs and 6.5 per 100 PYs respectively. Most patients who were changed from d4T-based regimens had symptoms of lipodystrophy, peripheral neuropathy, pain and neurological problems. In addition, majority of individual drug substitutions were done in patients on d4T-based regimens.

The majority of the patients were initiated on TDF-based (51%) regimens followed by d4T-based (35%). The overall burden of ART regimen changing was relatively lower than rates that have been reported in other studies. This might be due to the introduction of combined fixed-dose combination drugs that are less toxic and more potent [2, 8, 14, 19]. A study conducted by Cotte et al, reported ART modifications to be higher in patients on daily multiple drug regimens than those on FDC [23]. A study conducted in South Africa by Hirasen et al, reported that reducing pill burden or simplifying ART, improves adherence. Improvement in treatment adherence may lengthen first-line treatment, prevent treatment failure and ART changing to more expensive and limited second or third line therapy [24]. However, not all the patients qualify to take FDC therapy [10]. Patients with renal disease and drug resistance are excluded from the FDC regimens [10]. Overall, TDF-based regimens had a lower changing rate compared to d4T, AZT and non-standardised regimens.

The majority of patients (63%) were changed from d4T-based regimens. The changes were primarily due to ADRs, drug related toxicity and treatment failure. The higher rate of changing d4T-based regimen was consistent with the findings of the other studies conducted in both high and low income countries. The study conducted in a Regional Hospital, Kwa-Zulu Natal, reported d4T-based regimens having the highest frequency of changing ART regimens attributable to ADRs (65%) [20]. The other study conducted by Njuguna et al, reported TDF as the best tolerated drug with lesser drug substitutions (7.7%) as compared to d4T (46%) and AZT (27%) [22]. Another study conducted in Nairobi, Kenya, among sex workers, reported d4T-based (64%) being the most changed regimens as compared to TDF and AZT based [25].

The overall incident rate of ART regimen changes in the current study was comparable with the incident rate reported in Nairobi (11.1 per 100 PYs) [25]. The current study reports the overall incident rate of 10.0 per 100 PYs. The exposure time for patients initiated on non-standardised regimens was lower than all other regimens. The specific incident rates of regimen changes for d4T, AZT and TDF base regimens were 13.0, 8.0 and 6.5 respectively. We observed that TDF was more tolerated than d4T and AZT. The significant difference in changing rates were further observed on the KM plot that indicated the higher rate of changing the initial ART regimens in stavudine containing regimens followed by AZT and non-standardised regimens. These results are similar to the findings from the large cohort followed up for 10 years in Cape Town, South Africa [22]. These findings were also comparable with the study conducted in Nairobi [25].

With the improvement in ART over years, the reasons for regimen modification changed from merely treatment failure and drug resistance to several factors [Günthard]. The regimens can be changed for convenience, tolerability, simplification of treatment, drug interactions, pregnancy, treatment failure and drug toxicities [1, 3, 12, 14, 17]. In this study, initial ART regimens were mostly changed due to ADRs, prescriber’s decision, drug toxicity and treatment failure. Other factors included pregnancy, guideline related, chronic illnesses, unavailability of treatment and the use of other concomitant drugs.

The study conducted in Kenya also found a gradual increase in ART changes attributable to ADRs and treatment failure [25]. In April 2013, South Africa started rolling out the fixed-dose combination (FDC) pill, which might be the contributing factor for high rate of changes due to prescriber’s decision and guideline related [26]. FDC was introduced to improve adherence to treatment, although there were several more factors contributing to non-adherence [2, 10, 27]. In the current study, patients who changed from TDF-based regimens, were mostly changed from non-combined TDF-based regimens to FDC.

Lipodystrophy, peripheral neuropathy, pain and rash were the most commonly experienced ADRs in the current study. We also found higher frequency of ADRs in females 71% and d4T-based regimens causing more ADRs in female than males. These findings are comparable with the results from the study contacted in Tshepang, South Africa, with neuropathy, rash and lipodystrophy being the highly prevalent ADRs [13]. The study also found females being at higher risk of developing ADRs than males. Another study in South Africa found higher number of females having experienced ADRs in addition to higher number of female patients receiving ART [28]. The study also found peripheral neuropathy, lipodystrophy and skin reactions (rash) as the most commonly reported ADRs.

There are other studies that found ADRs playing the major role in ART changing [29-30]. In a multivariate analysis, there were increased hazard ratios for regimen changes related to d4T, AZT and non-standardised regimens. These findings are in line with other previous studies that have reported that d4T is one of the most intolerable drugs, and TDF as the most tolerable [22, 25]. The findings also showed increased hazards for regimen changes for ADRs, drug toxicity and treatment failure. The other significant predictor of ART regimen changes was CD4 cell count of 200 and more cells/mm3. Not many studies support this finding, however, the study conducted in Nairobi, Kenya among sex workers, reported an increased risk of ART regimen changing associated with cd4 count of ≥ 200 cells/mm3. The increased hazard was reported from the univariate analysis (HR 4.51; 95% CI: 3.65.6) [25]. The findings of this study may not be generalizable to all patients on HIV treatment because the study was conducted in only one pharmacovigilance sentinel site, in one of nine provinces in SA. The study was also restricted to first incident of ART regimen changing, which may have underestimated the true rates of regimen changes. Data on TDF-based regimens are available for six years as compared to 14 years of d4T and AZT based regimens, this may also have underestimated regimen changes related to the TDF-containing regimens. The study also did not grade the severity of ADRs and some ADRs might have been mistaken with advanced stage of HIV or other co-morbidities. However, the study was done using a robust dataset, with adequate sample size, long follow up period and minimal missing data in the descriptive characteristics.

## Conclusion

Several predictors of initial ART regimen modification for backbone NRTIs were investigated. The study reports reasonable incidence rate of initial ART regimen changes among HIV-infected patients in Witbank, South Africa. Patients changed their initial ART regimens mainly due to adverse drug reactions. Drug toxicity, type of backbone Nucleoside Reverse Transcriptase Inhibitors, CD4 count of 200 and more cells/mm3 and treatment failure were other predictors of initial ART regimen changing. Use of stavudine-based regimens was associated with increased risk of ART regimen changing due to lipodystrophy, peripheral neuropathy and pain. We recommend that clinicians should consider changing all patients who are still on stavudine-based regimens to avoid long term drug toxicities. We recommend further studies to identify predictors of stavudine toxicity because not all the patients on stavudine-based regimens experience toxicity.

## Acknowledgements

I would like to convey my sincere gratitude to the following: Lazarus Rugare Kuonza, Alfred Musekiwa, Robert Summers for co-authoring this paper. We would like to thank Pharmacovigilance Research Centre and its staff for allowing the researchers to access the database and their contribution; South African Field Epidemiology Training Programme staff and University of Pretoria for their inputs and authorization of the study. We appreciate the scientific writing advice provided by Dorothy L. Southern.

